# Evaluation of potential serum biomarkers for individuals at risk of multiple sclerosis

**DOI:** 10.64898/2026.04.25.715317

**Authors:** Kristin Mounts, YunDuo Liu, Masashi Fujita, Juliana Oyegunle, Tradite Neziraj, Susan V. Pollak, Renu Nandakumar, Nyater Ngouth, Sonya U. Steele, Irene Cortese, Charles C. White, Steven Jacobson, Daniel S. Reich, Philip L. De Jager

## Abstract

Circulating proteins have been widely investigated as potential biomarkers in multiple sclerosis (MS), yet findings across studies are often inconsistent, likely reflecting differences in disease stage, treatment exposure, and cohort composition. Studying individuals at elevated risk of MS prior to disease onset offers a unique opportunity to identify immune alterations that precede clinical disease while minimizing confounders.

Here, we investigated whether alterations in six previously MS-associated biomarkers are detectable and associate to underlying genetic susceptibility in two independent sample collections comprising people with MS (pwMS), healthy controls, and asymptomatic first-degree relatives of pwMS from the Genes & Environment in MS (GEMS) study cohort. The panel, representing complementary axes of MS immunopathology, included granzyme A (GZMA), MER tyrosine kinase (MERTK), interleukin-2 receptor alpha (IL2RA), osteopontin (SPP1), CD30 (TNFRSF8), and chitinase-3-like protein 1 (CHI3L1).

None of the proteins demonstrated associations with MS. A composite score constructed from externally derived effect estimates was not associated with MS status in either collection or in meta-analysis. Among asymptomatic first-degree relatives, the composite score was not significantly associated with group status. In contrast, an inverse correlation between SPP1 and the MS genetic risk score among GEMS participants was found (β = -0.246, p = 0.001).

Together, these findings suggest that several circulating proteins recently proposed as MS biomarkers are not robust tools to distinguish MS from healthy individuals. However, SPP1 levels are highlighted for further evaluation among at-risk individuals, and further work is needed to determine whether circulating immune signatures can capture the earliest stages of MS in at-risk individuals.

## Introduction

Multiple sclerosis (MS) is a chronic inflammatory and neurodegenerative disease of the central nervous system (CNS). Increasing evidence suggests that MS has a prolonged presymptomatic phase during which pathological changes precede clinical diagnosis by years. For instance, archival samples have uncovered evidence that neurofilament light chain (NfL), a marker of neuroaxonal injury, may be elevated several years before symptom onset^1,2^.

Circulating proteins have been widely investigated as potential diagnostic, prognostic, and monitoring biomarkers in MS^3–6^. Yet, there are currently no existing tools to deploy in individuals at risk of MS in an attempt to detect the earliest phase of the disease and to discover immune pathways that may contribute to disease susceptibility^7,8^. As a result, we evaluated serum analytes that have been previously associated with MS, hypothesizing that they may be altered in this earliest phase of the disease as well as after the onset of symptoms. Therefore, we included asymptomatic first-degree relatives of persons with MS (pwMS) from the Genes & Environment in MS (GEMS) study, who have an increased genetic susceptibility for and increased incidence of MS relative to the general population^9,10^.

We assembled a targeted serum biomarker panel representing complementary axes of MS immunopathology^11–16^, including peripheral immune activation (IL-2 receptor alpha [IL2RA]), immune regulation and Th2-associated signaling (CD30 [TNFRSF8]), cytotoxic effector function (granzyme A [GZMA]), innate immune clearance and repair pathways (MER tyrosine kinase [MERTK]), and markers linked to neuroinflammatory and glial responses (chitinase-3-like protein 1 [CHI3L1] and osteopontin [SPP1]). We investigated these circulating immune proteins using multiplex immunoassays across MS participants, healthy control participants, and GEMS participants from two independent sample collections to determine whether the selected immune biomarkers demonstrate reproducible associations with MS or enhanced genetic risk of MS.

## Results

### Descriptive characteristics

To conduct our study, we accessed serum samples from two collections: Columbia University Irving Medical Center (CUIMC) and the National Institute of Neurological Disorders and Stroke (NINDS). Participants included individuals at risk of MS from the GEMS study (n = 73 from CUIMC, n = 50 from NINDS), persons with MS (pwMS) (n = 50 from CUIMC, n = 53 from NINDS), and control participants (n = 53 from CUIMC, n = 94 from NINDS). Demographic characteristics are shown in **Table 1**. Within both sample collections, pwMS were older than control participants (p < 0.01 for all comparisons). In the NINDS collection, pwMS were also older than GEMS, and GEMS were older than HC (p < 0.01 for all comparisons). In the CUIMC collection, BMI and sex differed nominally between GEMS and control participants (p < 0.05 for both comparisons), while pwMS were similar to HC in these measures. No significant differences in BMI or sex were observed within the NINDS collection between pwMS and control participants or between GEMS and control participants (p > 0.05 for all comparisons). We note that the CUIMC GEMS samples were collected nationally using overnight room temperature shipping, while the NINDS GEMS participants were flown to NINDS as part of an earlier study^9,10^; those blood samples were collected and processed at NINDS as part of a structured research visit.

**Table 1.** Demographic characteristics of persons with MS (pwMS), healthy control participants (HC) and first-degree relatives (GEMS) in the CUIMC and NINDS sample collections. \**P* values represent between-collection comparisons (CUIMC vs NINDS) using t-tests for continuous variables and χ² tests for categorical variables. Disease duration was compared between MS groups using the Wilcoxon rank-sum test. **Age was significantly different (p < 0.01 for all comparisons) between pwMS and HC within both CUIMC and NINDS collections. Age was significantly different between GEMS and pwMS or HC in the NINDS collection (p < 0.01 for both comparisons), but not the CUIMC collection. Sex and BMI differences were observed only between CUIMC GEMS and HC (p < 0.05 for both comparisons). Values are presented as mean ± standard deviation for continuous variables (age) and median (IQR) for disease duration, and number (percentage) for categorical variables.

| Variable | CUIMC HC<br>(n = 53) | CUIMC pwMS<br>(n = 50) | CUIMC GEMS<br>(n = 73) | NINDS HC<br>(n = 94) | NINDS pwMS<br>(n = 53) | NINDS GEMS<br>(n = 50) | P value* |
| --- | --- | --- | --- | --- | --- | --- | --- |
| **Age\*\***, mean ± SD | 31.8 ± 9.7 | 39.7 ± 12.9 | 33.7 ± 9.6 | 40.0 ± 13.2 | 48.5 ± 14.6 | 35.8 ± 8.5 | **<0.001** |
| **BMI\*\*** category, n (%) |  |  |  |  |  |  | **0.016** |
| 18.5–30 | 43 (81.1%) | 27 (54.0%) | 69 (94.5%) | 65 (69.1%) | 43 (81.1%) | 41 (82.0%) |  |
| <18.5 or >30 | 10 (18.9%) | 23 (46.0%) | 4 (5.5%) | 29 (30.9%) | 10 (18.9%) | 9 (18.0%) |  |
| **Sex\*\***, n (%) |  |  |  |  |  |  | **<0.001** |
| Female | 37 (69.8%) | 34 (68.0%) | 63 (86.3%) | 49 (52.1%) | 32 (60.4%) | 30 (60.0%) |  |
| Male | 16 (30.2%) | 16 (32.0%) | 10 (13.7%) | 45 (47.9%) | 21 (39.6%) | 20 (40.0%) |  |
| **Race**, n (%) |  |  |  |  |  |  | 0.46 |
| African American | 11 (20.8%) | 8 (16.0%) | 3 (4.1%) | 15 (16.0%) | 8 (15.1%) | 0 (0%) |  |
| Asian | 3 (5.7%) | 0 (0%) | 1 (1.4%) | 3 (3.2%) | 5 (9.4%) | 0 (0%) |  |
| Caucasian | 35 (66.0%) | 34 (68.0%) | 69 (94.5%) | 67 (71.3%) | 38 (71.7%) | 50 (100%) |  |
| Other / Hispanic | 4 (7.5%) | 8 (16.0%) | 0 (0%) | 9 (9.6%) | 2 (3.8%) | 0 (0%) |  |
| **Disease duration (years)**, median (IQR) | — | 0.0 (0.0–0.2) | — | — | 7.6 (2.4–11.9) | — | **<0.001** |

We also note that NINDS pwMS had substantially longer disease duration than in CUIMC (p < 0.001), consistent with the fact that CUIMC pwMS were sampled during their diagnostic evaluation whereas NINDS pwMS were collected cross-sectionally, often later in the disease course. Disease subtype and immunomodulatory treatment characteristics are presented in **Supplementary Table 1**. CUIMC pwMS included untreated (n = 32) and treated (n = 18) participants. NINDS pwMS included treatment naïve (n = 50) and treated participants (n = 3).

### Analyte selection strategy

We assembled a targeted panel of circulating immune proteins that had previously been reported to be altered in pwMS relative to control participants or in disease course (**Table 2**, **Figure 1**). PubMed-indexed studies were selected using search terms including “multiple sclerosis,” “biomarker,” “serum,” “plasma,” “cerebrospinal fluid.” We prioritized analytes with associations across multiple studies and measured using comparable immunoassay platforms. CHI3L1 and SPP1 were selected based on extensive prior evidence linking them to glial activation and neuroinflammatory signaling in MS, with elevations reported in both CSF and blood, and strong performance in multivariate biomarker models^11,13,17–19^. IL2RA and TNFRSF8 were incorporated as circulating indicators of T-cell activation and immune regulation associated with disease activity and early disease states, respectively^14–16,19^. To incorporate signals from recent large-scale analyses, GZMA and MERTK were included based on findings from a plasma proteomics study in the UK Biobank resource; this study reported decreased GZMA and increased MERTK levels in MS^12^, implicating alterations in cytotoxic lymphocyte function and monocyte-microglial signaling pathways. Together, this panel was designed to capture complementary axes of MS biology that may be altered early on in genetically at-risk individuals, including peripheral immune activation and downstream neuroinflammatory processes.

**Figure 1.**
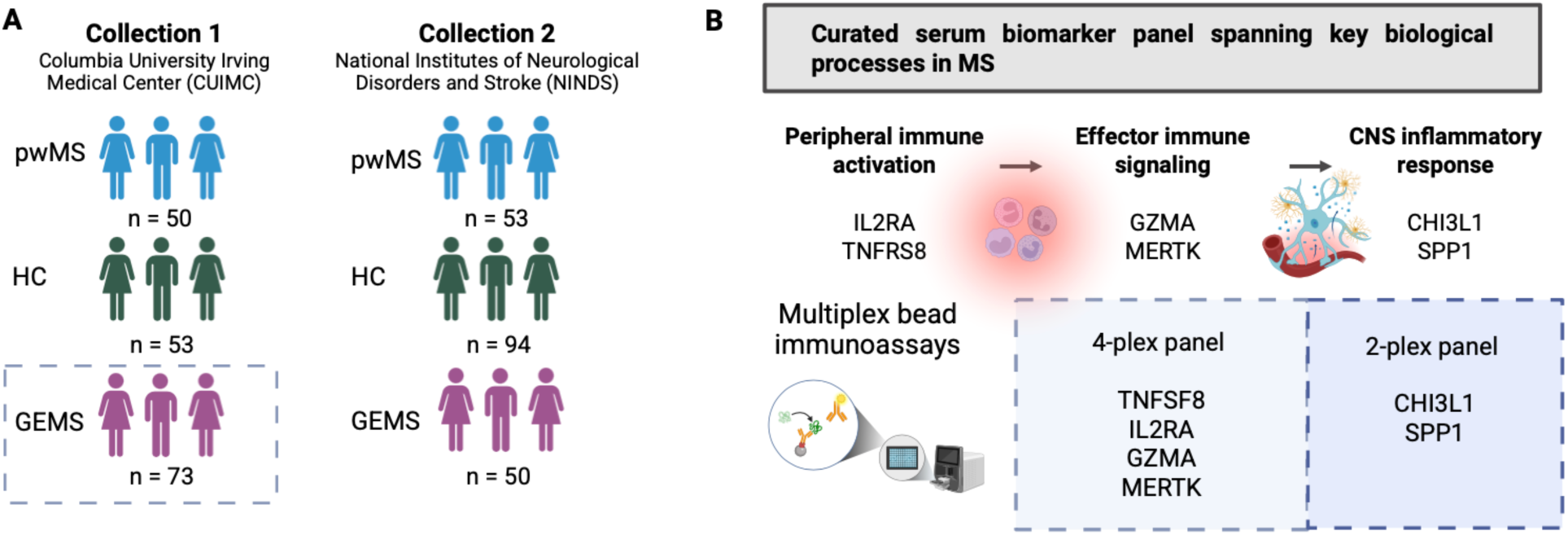
Study sample collections and biomarker panel design. **(A)** Serum samples were obtained from two independent sample collections: Columbia University Irving Medical Center (CUIMC) and the National Institute of Neurologic Disorders and Stroke (NINDS), each included persons with multiple sclerosis (pwMS), healthy controls (HC), and asymptomatic first-degree relatives from the Genes and Environment in Multiple Sclerosis study. The GEMS samples obtained at NINDS were processed the same day, using the same procedure as the other sample classes. The dotted box around the CUIMC GEMS participants denotes the fact that these samples were collected nationally and shipped overnight to Columbia where they were processed using the same procedure as the other sample classes. **(B)** Six different proteins previously evaluated in the context of MS were prioritized for evaluation in our study. They capture different immune responses relevant to MS. Given their concentration in peripheral blood, two different multiplex assay panels were developed.

**Table 2.**
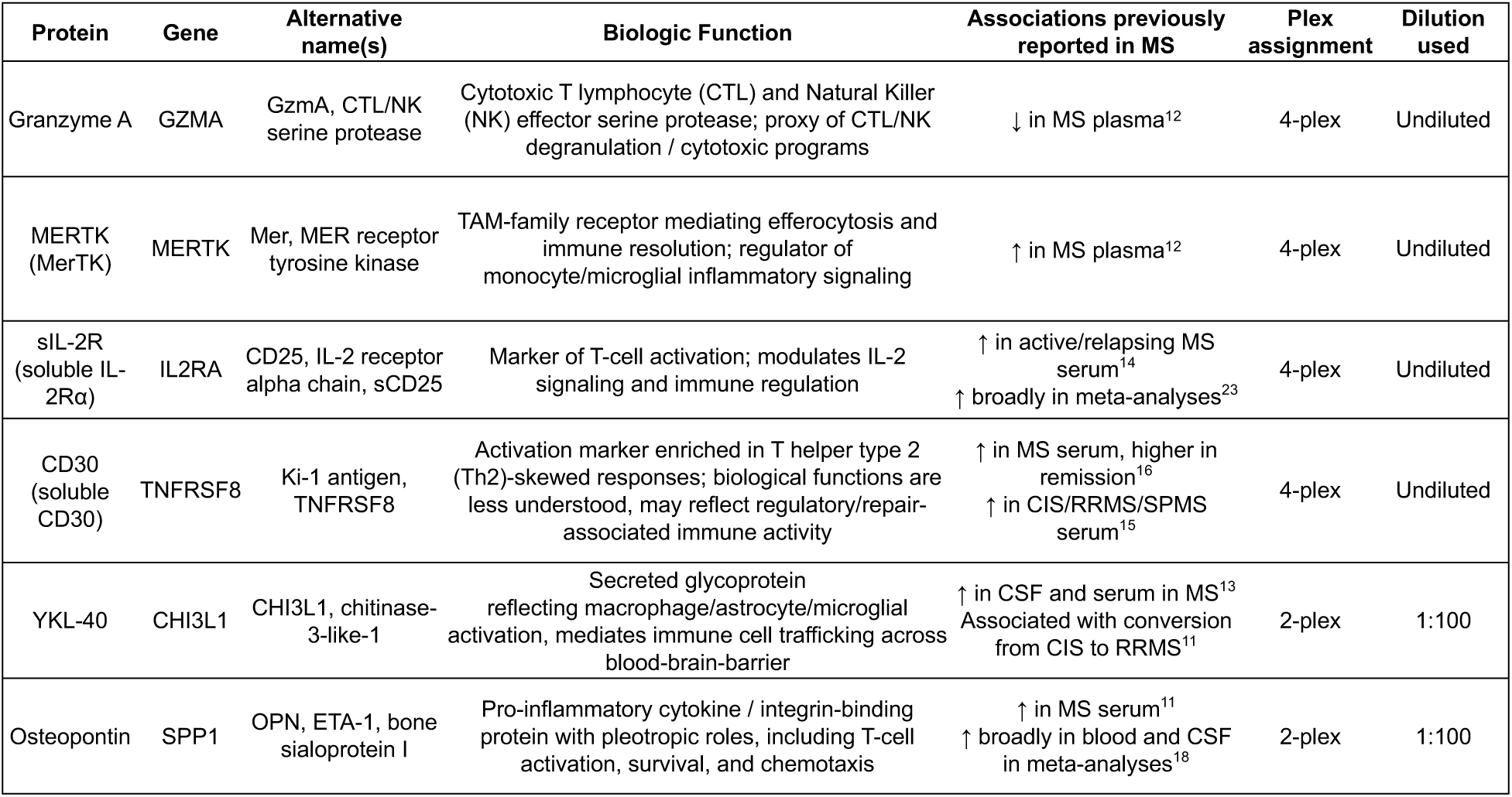
Curated serum biomarker panel and assay design. Summary of the six proteins included in the multiplex immunoassay panel used in this study. For each analyte, the table lists the corresponding gene symbol, commonly used alternative names, major biologic functions, and previously reported directions of association with multiple sclerosis (MS) based on prior literature. Proteins were selected to represent complementary aspects of MS immunobiology shown here. The table also indicates the multiplex assay configuration (plex assignment) and dilution used for each analyte in the Procartaplex bead-based immunoassay. Analytes were measured either within a four-analyte panel (4-plex) or a two-analyte panel (2-plex) according to assay optimization requirements. Directions previously reported in MS refer to associations observed in serum, plasma, or cerebrospinal fluid across the highlighted studies and meta-analyses.

### Circulating proteins and a composite score are not associated with MS or with at-risk individuals

We first attempted to replicate the reported associations of each analyte (GZMA, MERTK, IL2RA, SPP1, TNFRSF8, and CHI3L1) with a diagnosis of MS using linear regression models adjusted for age, sex, BMI, and batch within each dataset. In both sample collections, there were no significant associations (**Table 3** and **Supplementary Figure 1**). To assemble all of the evidence for each analyte, we deployed a random-effects meta-analysis with restricted maximum likelihood (REML) estimation of between-collection variance. None of the six analytes demonstrated significance in these analyses (**Table 3**, **Figure 2a**). For most analytes, the direction of effect was consistent in the two sample collections, although the magnitude of these effects was small and not statistically significant in the meta-analysis. We do note, in both sample collections, the expected association of greater serum CHI3L1 with advancing age at a nominal p < 0.05 threshold of significance (CUIMC: β = 0.014, p = 0.022, FDR = 0.11; NINDS: β = 0.012, p = 0.011, FDR = 0.12) and in meta-analysis (β = 0.013, p < 0.001), consistent with a prior report^20^. No analytes were associated with BMI or female sex after multiple testing correction (**Table 3**).

**Figure 2.**
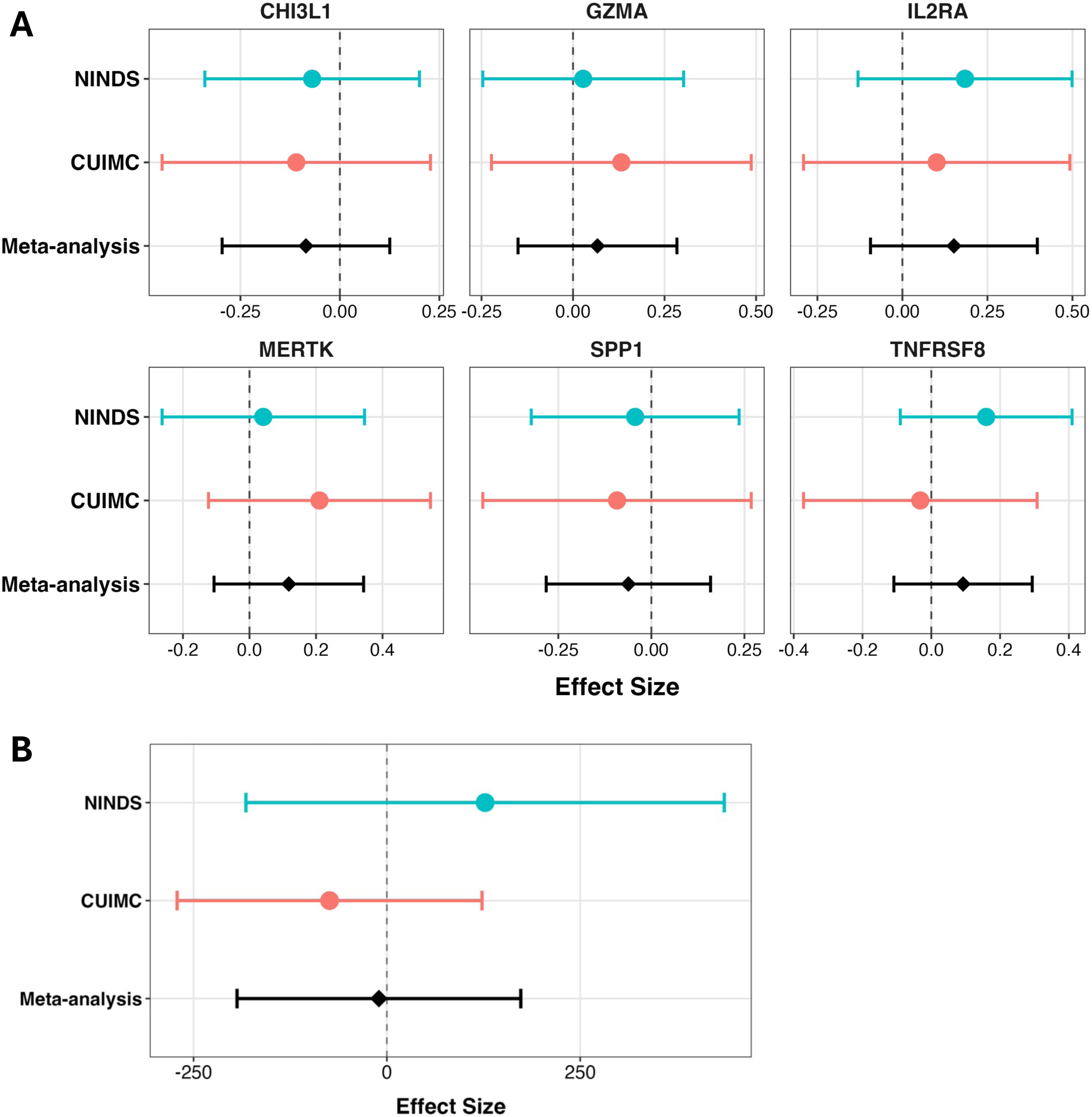
Lack of association of individual serum proteins or the derived composite protein score with MS. **(A)** Forest plots show collection-specific and the meta-analysis estimates for case-control comparisons based on rank-inverse normal transformed (RINT)-transformed protein levels. Points represent effect estimates with 95% confidence intervals (CIs). Pooled estimates were calculated using random-effects meta-analysis with restricted maximum likelihood (REML) estimation of between-collection variance. The dashed vertical line indicates the null effect (β = 0). **(B)** Collection-specific and the meta-analysis estimates for the association between the composite protein score and MS status.

**Table 3.**
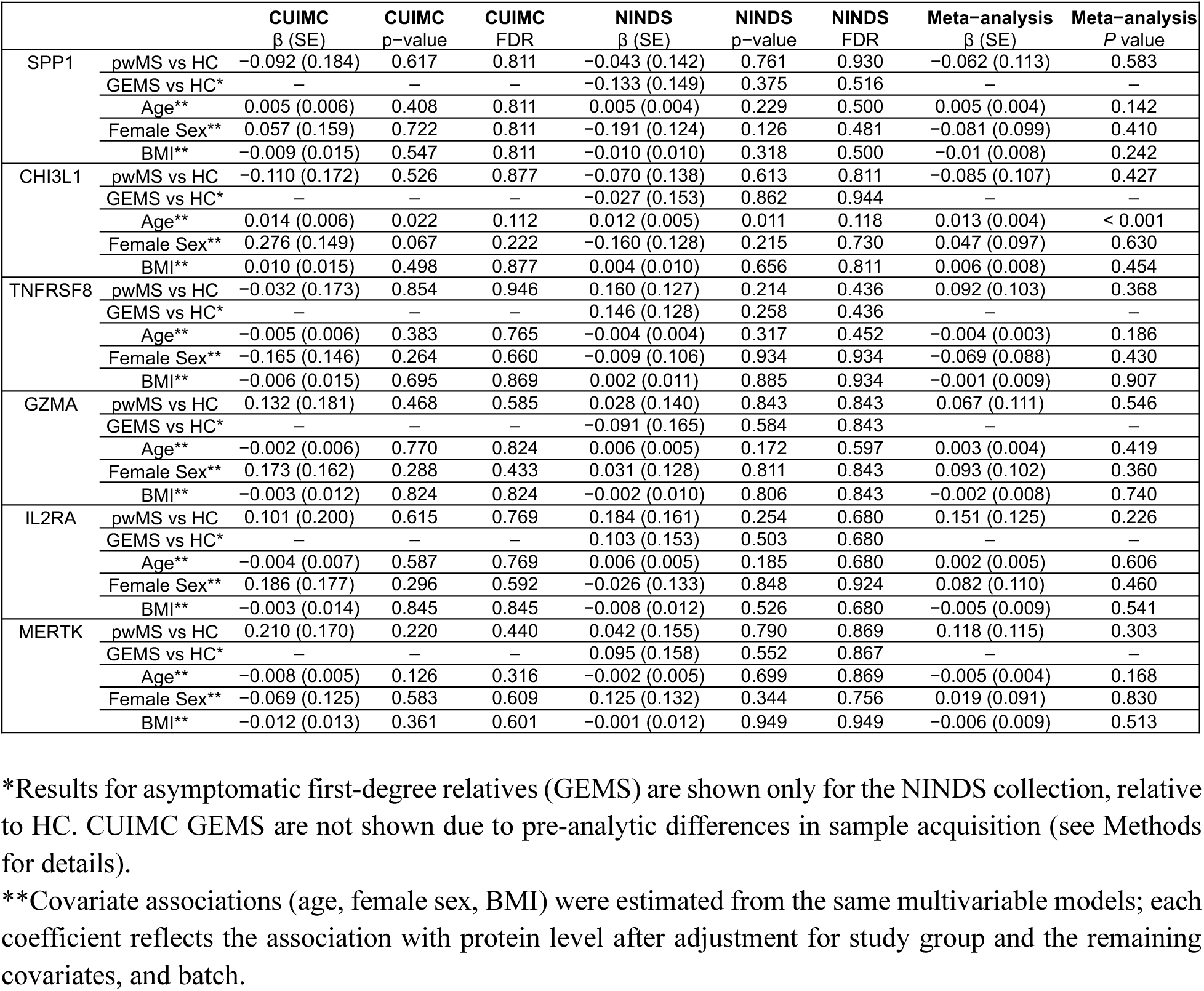
Associations of individual serum proteins with MS status, first-degree relative status, and covariates. Linear regression models were fit separately in CUIMC and NINDS using rank inverse normal transformed (RINT) protein levels as outcomes. Models included study group (Subject_Type: pwMS, HC, GEMS) as a categorical predictor and were adjusted for age, sex, BMI, and batch. A summary of the evidence for MS case–control comparisons was obtained by combining collection-specific regression coefficients and standard errors using random-effects meta-analysis with restricted maximum likelihood (REML). Results are presented as regression coefficients (β) with standard errors (SE), nominal p-values, and false discovery rate (FDR)–adjusted p-values.

We then compared the analytes between the GEMS and control participants. We note that the CUIMC GEMS samples were collected nationally, using overnight shipping from the participant’s location, while all other sample categories (including the NINDS GEMS samples) were collected at one site and processed the same day. We therefore limit these analyses to the NINDS dataset. Among the NINDS participants, for whom all samples were collected and processed under uniform conditions, we found no significant differences between the GEMS participants and either control participants or pwMS (**Table 3**).

Because individual analytes often show modest effect sizes and analyte measurements can be noisy, we next tested whether a multi-protein signature could be an alternative approach to assess the utility of the selected analytes. Here, we constructed a weighted composite score using previously published regression effect estimates as weights^12^. In regression models adjusted for covariates, the composite score was not significantly associated with MS diagnosis in either sample collection (**Supplementary Table 2**), and the meta-analysis showed no significant association between the composite protein score and MS status (**Figure 2b**) (pooled β = -5.39, SE = 95.12, 95% CI -191.81 to 181.04, p = 0.955). Thus, the aggregate proteomic signal did not reproducibly distinguish pwMS from healthy controls in this dataset. Cochran’s Q test did not detect statistically significant heterogeneity across the two sample collections (Q = 1.09, p = 0.296). The estimated between-study variance was τ² = 1680.82 and I² = 8.3%.

For GEMS participants, this comparison was performed in the NINDS collection only, where the composite score showed a positive but non-significant association relative to controls (β = 251.026, p = 0.121, **Supplementary Table 2**). No significant associations were observed with age, female sex, or BMI in either collection (**Supplementary Table 2**).

### Genetic risk associations with serum proteins and composite score in first-degree relatives

Because first-degree relatives represent a genetically enriched population for risk-associated alleles at MS susceptibility variants, we next examined whether circulating analytes were associated with summary measures of genetic risk within this group. Genetic risk scores (GRS) were calculated from genotyping data using prior established weighted sums of MS-associated susceptibility variants^9,10^. We tested versions with and without major histocompatibility complex (MHC) loci as well as an MHC-only score (**Supplementary Table 3**, **Supplementary Figures 2-4**).

These analyses were run in each of the datasets and combined using a meta-analysis approach (**Table 4**, **Figure 3a**). This meta-analysis revealed a significant association of SPP1 with the GRS64 score (β = -0.24, p = 0.0013), indicating that higher genetic risk for MS was associated with lower circulating SPP1 levels. No other analytes showed statistically significant associations with genetic susceptibility in the meta-analyses (**Table 4**). This effect was observed in each of the two component datasets, and the direction was consistent in both (decrease). However, only the NINDS dataset was significant on its own.

**Figure 3.**
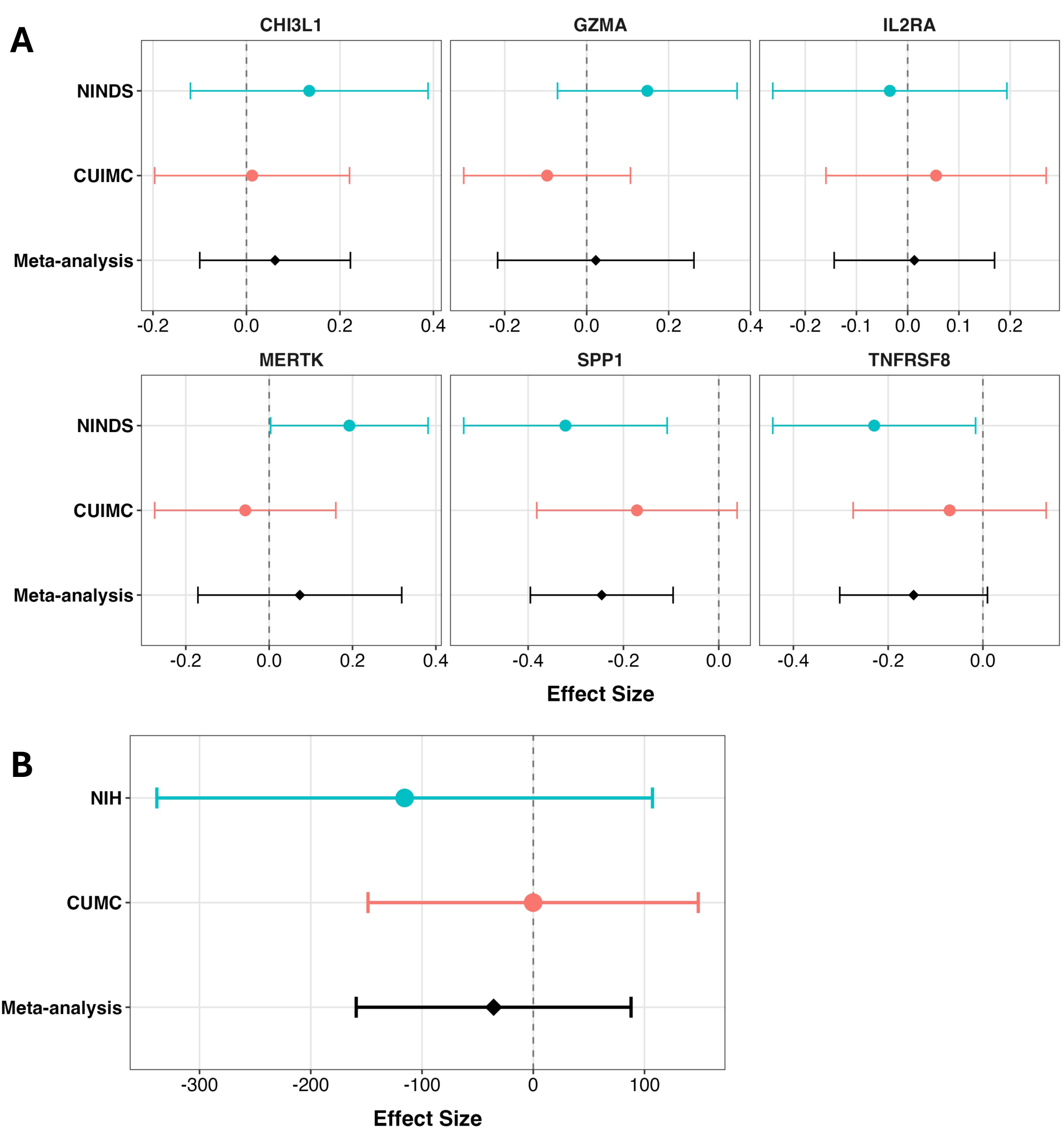
Associations of MS genetic risk with circulating proteins and the composite protein score in first-degree relatives. Forest plots summarizing associations between the MS genetic risk score (GRS64) and circulating immune markers in GEMS participants from the CUIMC and NINDS collections. Within each collection, linear regression models were fit with rank-inverse normal transformed (RINT) protein levels as outcomes and GRS64 as the primary predictor, adjusting for age, sex, BMI, and batch. Points represent collection-specific β estimates with 95% confidence intervals (CIs). A random-effects meta-analysis with restricted maximum likelihood (REML) estimation of between-collection variance was used to provide a summary of the evidence. The dashed vertical line indicates the null effect (β = 0). **(A)** Meta-analysis of associations between GRS64 and individual serum proteins. **(B)** Meta-analysis of the association between GRS64 and the weighted composite protein score.

**Table 4.**
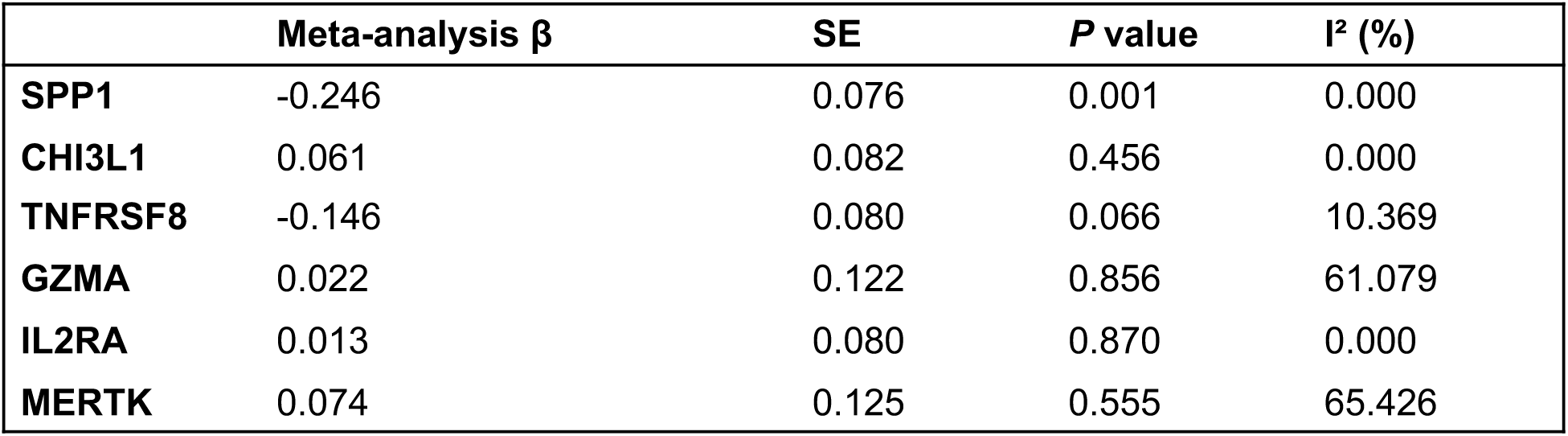
Meta-analysis of associations between circulating protein levels and MS genetic risk (GRS64). Within each collection, linear regression models were fit with RINT-transformed protein levels as outcomes and GRS64 as the primary predictor, adjusting for age, sex, BMI and batch. Collection-specific regression coefficients (β) and standard errors (SE) were combined using random-effects meta-analysis with restricted maximum likelihood (REML). The meta-analysis β represents the pooled regression coefficient describing the change in protein level per unit increase in GRS64. *P* values correspond to the effect estimate. Between-collection heterogeneity was assessed using the I² statistic.

A notable suggestive result is seen with MERTK which showed a positive association with total GRS in the NINDS dataset (r = 0.419, p = 0.005, FDR = 0.031); there was a slightly stronger association with the MHC-only GRS (r = 0.463, p = 0.002, FDR = 0.011) than the non-MHC GRS (r = 0.236, p = 0.128, FDR = 0.383). In the meta-analysis, the result was not significant; larger sample sizes will be needed to explore this observation further.

Finally, we evaluated whether genetic risk was associated with the composite protein signal. Regression models testing the relationship between the composite score and GRS64 within each collection showed no association in the meta-analysis of both sample collections (**Figure 3b**; β = -35.6, SE = 63.0, 95% CI -159.1 to 87.9, p = 0.571).

## Discussion

In this study, we did not replicate the association of the prioritized candidate serum biomarkers with a diagnosis of MS, and we did not find a significant difference between GEMS and healthy participants for these markers. Similarly, aggregating the effects of the 6 markers into a single score did not yield a significant difference between healthy participants and either MS or GEMS participants. However, we did find an intriguing association between the MS GRS and serum SPP1 levels.

The development of reliable MS biomarkers is challenged by limited replication of single-protein findings, the biological heterogeneity of MS, and diversity of patient collections that have been used in different studies. Reported associations for circulating proteins, including those included in our panel, vary with disease course, treatment exposure, as well as age and weight^11,13–30^. Many markers, including those in our panel, have also been reported across other autoimmune^31–37^ and inflammatory conditions^16,18^, suggesting that they primarily reflect non-specific inflammatory responses rather than MS-specific pathology.

These findings align with a recent UK Biobank-based plasma proteomics study in MS, where most protein associations overlapped with other immune-mediated conditions^12^. Other previously MS- associated markers, including several of those tested here, also failed to be replicated in that analysis^12,38^. Moreover, this large study reported limited replication of their own results in external cohorts profiled using the same platform: most prioritized analytes in the discovery analysis only demonstrated directional concordance of an analyte’s non-significant effect^12,38^. These results suggest that, while certain analytes may show reproducible associations with MS, the effect sizes are small and unlikely to yield a very informative clinical biomarker unless its context is well constrained.

Our lack of replication of prior observations could therefore simply be due to limited statistical power to reproduce the association of analytes of modest effect sizes. Our sample size was moderate relative to the earlier large population proteomic studies: the minimum detectable effect size at 80% power was a Cohen’s d ≈ 0.36. Consistent with the large UK Biobank-based study, we therefore confirm that our target analytes do not have a strong effect in terms of an association with MS, and their potential for utility in a clinical setting may therefore be limited. Additional limitations include the measurement variability in the selected analytes across platforms^39,40^ and the biofluid itself, both of which could influence the detection of subtle disease associations, particularly for low-abundance immune markers where plasma appears to be the more sensitive substrate^41,42^.

There were also notable differences between the two collections of samples since we are repurposing samples collected as part of earlier studies. First, the distribution of genetic risk scores differed between the two collections: the NINDS GEMS participants had been selected using the MS GRS, focusing on the top and bottom 15% of GEMS GRS scores. On the other hand, the CUIMC GEMS participants were sampled from the entire GRS distribution. The NINDS samples may exaggerate an association that is driven by outliers in MS GRS. Second, there were differences in demographic features between participant groups, given constraints in sample availability, and we therefore accounted for these differences by adding covariates to our equations. The pwMS were also distinguished by individuals early in their disease course (CUIMC) and individuals sampled throughout their disease course (NINDS). The larger proportion of treated individuals in CUIMC relative to NINDS pwMS may also skew results. Finally, the difference in sample acquisition between the CUIMC GEMS (blood shipped overnight) and the pwMS and the control participants (blood processed on day of collection) led to a greater variance in analyte measures (**Supplementary Figure 1**).

Despite these limitations, we uncovered a significant result with SPP1 and the MS GRS (**Figure 3**). SPP1 has been shown to promote pro-inflammatory T cell responses and contribute to neuroinflammation^43,44^, and circulating levels reflect both genetic and inflammatory influences^45^. The observed inverse relationship in asymptomatic first-degree relatives may reflect a distinct immune state that contributes to disease onset. Alternatively, as the average age of our GEMS participants (**Table 1**) is slightly greater than the typical average age at symptom onset for pwMS, individuals that are resilient to factors influencing the onset of MS may be somewhat enriched in our collection, and the reduced SPP1 level may reflect an immune set point associated with resilience or delayed disease onset. A non-mutually exclusive explanation is that early inflammatory activity may be compartmentalized within the CNS, as SPP1 is highly expressed within MS lesions^46^ and CSF levels may better reflect neuroinflammation in MS than circulating levels^47,48^. In this context, early neuroinflammatory activity associated with genetic susceptibility may not yet be reflected in peripheral circulation. However, these interpretations of reduced SPP1 serum levels in relation to higher MS GRS remain speculative, and validation of the observation in additional samples is necessary.

MERTK demonstrated a significant association with genetic risk in the NINDS GEMS participants, but this was not observed in the CUIMC collection and was not significant in the meta-analysis. This discrepancy may reflect the underlying differences in the distribution of genetic risk between CUIMC and NINDS GEMS participants, as the latter were enriched for individuals at the extremes of genetic risk and may have increased power to detect associations. Since the meta-analysis is not significant, we consider this result to be suggestive and worthy of further evaluation. The direction of effect is consistent with expectation of greater MERTK levels in MS.

In summary, several circulating proteins previously proposed as MS biomarkers did not demonstrate large or reproducible cross-sectional differences between MS and healthy individuals across independent sample collections. This is consistent with the fact that there are no clinical biomarker assays for MS diagnosis. While these proteins may have some association with MS, the effect sizes are generally quite modest and context specific. They may primarily reflect dynamic immune variation rather than stable disease identity and thus have limited utility as cross-sectional measures in individuals at risk of MS. Large prospective, longitudinal studies of at-risk individuals are needed to identify better biomarkers that could be used for early diagnosis and as outcome measures in trials. The significant relation of SPP1 levels to the MS GRS suggest that some circulating proteins may capture baseline immune states associated with inherited susceptibility. Here also, longitudinal studies will be required to determine how this immune measure evolves from peripheral immune changes to the earliest stages of MS in which the target organ, the CNS, is disrupted by an infiltrative inflammatory process.

## Materials and Methods

### Study collections

#### CUIMC pwMS and control participants

Serum samples from 50 pwMS and 53 healthy control participants from Columbia University Irving Medical Center (CUIMC) were analyzed (**Figure 1**). MS participants were approached and consented prior to undergoing a diagnostic lumbar puncture as part of the work-up to establish a diagnosis of MS. Blood was drawn after the procedure, processed to prepare serum and then cryopreserved at -80 °C until it was thawed for our assay. Demographic information, immunomodulatory treatment history, and MS disease status were obtained from patient charts (**Table 1** and **Supplementary Table 1**). Healthy control participants aged 18 years and older were recruited from the New York City metropolitan area using printed and electronic advertisements. Height, weight, and body mass index (BMI) were recorded at study visit. Demographic and medical history data was obtained using standardized questionnaires. Included participants reported no history of chronic infectious, inflammatory, and metabolic diseases.

#### GEMS first-degree family relatives

Samples from asymptomatic first-degree relatives of pwMS were drawn from the Genes and Environment in Multiple Sclerosis (GEMS) project, a nationwide prospective study of genetic and environmental determinants of MS risk, comprised of over 1,900 participants followed longitudinally to capture the transition from health to clinical MS^9^.

Samples were collected via two different avenues: (1) a nationwide effort using local phlebotomy and overnight shipping to a central processing site (CUIMC GEMS; n = 73); and (2) a sub-study in which selected GEMS participants traveled to NINDS for in-depth evaluation (NINDS GEMS; n = 50), during which a blood sample was obtained and processed. Samples from the NINDS participants are repurposed from our earlier study comparing the phenotypic characteristics of individuals with low or high Genetic and Environmental Risk Score for Multiple Sclerosis Susceptibility (GERS_MS_)^10^. Additional details regarding participant selection and risk score derivation have been described previously^10^. In contrast, CUIMC GEMS participants were not selected based on genetic risk; they represent a subsample of the study population.

#### NINDS collection

This collection consists of 53 pwMS and 94 control participants collected at the National Institute of Neurologic Disorders and Stroke (NINDS). The 50 pwMS samples were selected from the Natural History of Multiple Sclerosis protocol (NCT00001248), which includes participants who were treatment-naïve at the time of sample collection; an additional three pwMS samples were included from the NINDS GEMS sub-study. Healthy control participants were obtained from two sources: 50 samples from the National Institutes of Health Blood Bank (Department of Transfusion Medicine) and 45 samples were part of the NINDS GEMS sub-study. Because all samples underwent uniform collection and processing procedures on-site at NINDS, direct comparisons among pwMS, HC, and GEMS were performed within this sample collection.

At the two sites, all participants provided written informed consent, and all study procedures were approved by the Columbia or NINDS institutional review boards (IRB #AAAL4106, IRB #AAAT5674; NCT01617395, NCT00001248).

### Serum collection and processing

CUIMC pwMS and healthy control samples were collected in BD Vacutainer serum tubes (red-top, clot activator) and processed on-site. GEMS samples (CUIMC) were shipped as whole blood overnight at room temperature and processed into serum upon receipt. Following processing, serum samples were stored at -80 °C.

NINDS samples were collected, processed, and stored at -80 °C onsite and subsequently shipped to CUIMC overnight on dry ice. Blood samples from pwMS and GEMS were collected in serum separator tubes (SST), while HC samples were obtained from the blood bank using standard blood collection tubes.

All samples were centrifuged at 2000 rpm for 10 minutes at room temperature, aliquoted (1 mL) into cryovials and stored at -80 °C. The time from blood draw-to-freezing was between 3 and 4 hours. Samples were thawed for the first time at CUIMC immediately prior to the experiment.

### Multiplex bead-based immunoassays

Serum protein concentrations were measured using custom multiplex bead-based immunoassays (ProcartaPlex, Thermo Fisher Scientific). Samples were run in duplicate and processed in batches over a one-year period. Samples were randomized across plate positions and balanced across study groups to minimize batch effects. Fluorescence intensity was measured using a Luminex 200 instrument.

### Data pre-processing and quality control

Analyte concentrations were calculated using standard curves consisting of six or seven calibration points and fit using a five-parameter logistic regression model within the ProcartaPlex analysis software (ProcartaPlex Analysis App). Samples with values below the lower limit of quantification were replaced with one-half of the assay’s lower limit of detection for the analyte.

During data quality evaluation, we noted that CUIMC GEMS samples demonstrated significant differences in two analytes compared with healthy control participants: MERTK (β = 0.72, p = 4 × 10⁻⁶, FDR = 3.6 ×10⁻^5^) and CHI3L1 (β = 0.48, p = 0.0048, FDR = 0.048) levels were increased. However, because these samples were collected differently (shipped as whole blood overnight prior to serum isolation) whereas other samples were processed on-site shortly after collection, we considered the possibility that shipment or delayed handling could influence the protein levels. Consistent with this concern, the observed associations were not replicated in the NINDS first-degree relatives (**Table 3**), where sample collection and processing was uniform across groups. Therefore, these observations were interpreted cautiously and considered as potential artifacts due to the differences in sample collection.

Assay performance metrics, including intra- and inter-assay variability, are provided in **Supplementary Table 4**. Additional details regarding assay procedures and quality control are provided in **Supplementary Methods**.

### Statistical analysis

#### Case-control and at-risk comparison

Normality of serum protein concentrations was assessed using Shapiro-Wilk tests, and all six analytes deviated significantly from a normal distribution. Concentrations were rank-transformed and mapped to the standard normal distribution (rank-inverse normal transformation, RINT) prior to analysis (**Supplementary Figure 5**). Within each collection, RINT-transformed protein levels were modeled using linear regression with study group (Subject_Type) as a categorical predictor and adjustment for age, sex, BMI, and experimental batch/date. Covariate associations were estimated from the same collection-specific multivariable linear models; thus, each covariate coefficient reflects its association with the transformed protein level after adjustment for study group, the remaining covariates, and experimental date (batch). Multiple testing correction was performed using the Benjamini-Hochberg false discovery rate (FDR) procedure. All analyses were restricted to samples with non-missing values, and participants with missing or unknown age, sex, or BMI were excluded (n = 2, healthy controls).

### Meta-analysis

To summarize effects across the two sample collections, we combined collection-specific estimates using random-effects meta-analysis with restricted maximum likelihood (REML). Here, we used a multilevel random-effects model to account for the nested structure of collection-specific estimates within biomarkers, allowing estimation of the overall effect while accommodating between-biomarker and between-collection variability. For each analysis, regression coefficients and corresponding standard errors were first estimated separately within each collection using identical model specifications. These collection-specific estimates were then pooled to obtain overall effect estimates and confidence intervals. This framework was applied to: (1) case-control comparisons of individual biomarkers; (2) analyses evaluating the composite protein score in relation to MS status; (3) associations between circulating protein levels and MS genetic risk scores (GRS); and (4) associations between the composite protein score and GRS among GEMS participants. Between-collection heterogeneity was summarized using Cochran’s Q and I² where applicable.

### Composite score

To evaluate whether a multi-protein signature was present in our collections, we constructed a weighted composite protein score using externally derived regression coefficients^12^. For each protein, the RINT-transformed concentration was multiplied by the corresponding β coefficient, and the weighted values were summed across all proteins to generate a single aggregate score for each subject:

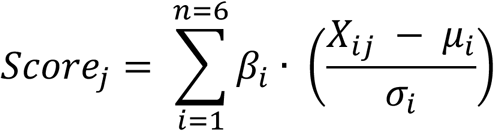

Where Score_j_ represents the composite score value for subject *j*, *β_i_* is the published regression coefficient (weight) for protein *i*, *X_ij_* is the RINT-transformed concentration for protein (i) for subject *j*, *μ_i_* and σ_i_ are the mean and standard deviation, respectively, of protein *i* in the collection, and the 6 biomarkers (*n*). The signs of the reported coefficients were preserved to maintain the biological direction of the associations.

Under this framework, higher composite scores represent individuals whose profiles more closely match the pattern reported in the prior study (specifically, higher levels of proteins positively associated with MS and lower levels of proteins negatively associated with MS). The composite score was analyzed as a continuous outcome in collection-specific linear regression models for group comparisons and covariate associations, with adjustment for age, sex, BMI, and batch/date, and in separate models evaluating association with genetic risk scores. Composite score comparisons across study groups were restricted to samples collected and processed under uniform conditions; CUIMC GEMS data were excluded.

### Association with genetic risk score

Genetic risk for MS among GEMS participants was quantified using previously validated, weighted genetic risk scores (GRS)^9,10^ derived from MS susceptibility variants identified in genome-wide association studies that were current at the time of recruitment, including large meta-analyses from the International Multiple Sclerosis Genetics Consortium (circa 2011)^49,50^. Three related GRS were evaluated: (1) the GRS64, which aggregates the effects of 64 MS-associated single nucleotide polymorphisms; (2) an MHC-specific GRS, which aggregates the effects of MS risk variants found within the MHC (includes HLA-DRB1*15:01, HLA-A*02:01, HLA-DRB1*04:01, HLA-DRB1*04:04, and HLA-DRB1*14:01) and (3) a non-MHC GRS, capturing the cumulative contribution of non-MHC susceptibility loci. Variants were coded additively and weighted by published effect sizes, and have been shown to stratify MS risk within the GEMS cohort^9,10^. A complete list of variants, genomic annotations, and corresponding weights used to construct these genetic risk scores is provided in **Supplementary Table 5**.

Within each collection, associations between RINT-transformed protein levels and GRS were first evaluated using Pearson correlation coefficients. For the primary analysis of GRS64, collection-specific linear regression models were fit with RINT-transformed protein levels as the outcome and GRS64 as the primary predictor, adjusting for age, sex, BMI, and batch/date. Collection-specific regression coefficients and standard errors were then combined using random-effects meta-analysis with REML. Associations of GRS64 with composite score were performed across all GEMS participants, including CUIMC.

A post hoc power calculation was performed to estimate the minimum detectable standardized effect size given the observed variance in the data, assuming a two-sided α = 0.05 and 80% power.

### Software and data availability

All analyses were performed using R (version 4.4.1). Additional details are provided in the **Supplementary Methods**. Analysis code will be made available upon publication. De-identified data supporting the findings of this study is available from the authors upon request and subject to institutional and ethical approvals.

## Supplementary Materials

**Supplementary Methods**

**Supplementary Figure 1.** Distribution of RINT-transformed serum protein levels

**Supplementary Figure 2.** Associations with GRS64

**Supplementary Figure 3.** Associations with MHC-only GRS

**Supplementary Figure 4.** Associations with non-MHC GRS

**Supplementary Figure 5.** RINT transformation distributions

**Supplementary Table 1.** Clinical characteristics

**Supplementary Table 2.** Composite score associations

**Supplementary Table 3.** GRS correlations

**Supplementary Table 4.** Assay performance

**Supplementary Table 5.** Genetic variants and weights

## Acknowledgements

We thank the individuals who participated in this study. Procartaplex assays were performed by the Biomarkers Core Laboratory at the Irving Institute for Clinical and Translational Research (Susan V. Pollak, Renu Nandakumar), under award UL1TR001873. This work was supported by National Multiple Sclerosis Society (RG-5003-A-2; P.L.D.J.). K.M. was supported by the National Institutes of Health Medical Scientist Training Program (T32GM145440). This research was also supported in part by the Intramural Research Program of the National Institutes of Health (NIH). The contributions of the NIH authors are considered Works of the United States Government. The findings and conclusions presented in this paper are those of the authors and do not necessarily reflect the views of the NIH or the U.S. Department of Health and Human Services. Figure 1 was created using BioRender (https://www.biorender.com).

## Author contributions

K.M., Y.L., and P.L.D. contributed to the study design and methodology; S.J., D.S.R., and P.L.D. sample collection; K.M., S.V.P. data acquisition; K.M., Y.L., C.C.W., M.F. statistical analysis; K.M., S.J., D.S.R., C.C.W., J.O., N.N., and P.L.D. data curation; K.M. and P.L.D. manuscript draft; K.M., Y.L., M.F., J.O., T.N., S.V.P, R.N., N.N., S.U.S., I.C., C.C.W, S.J., D.S.R., and P.L.D. manuscript, review and editing. P.L.D. obtained funding for the study.

## Conflict of Interest

The authors declare no conflicts of interest.

## Notes

### Competing Interest Statement

The authors have declared no competing interest.

