## Supplementary Figures and Tables for "Evaluation of potential serum biomarkers for individuals at risk of multiple sclerosis"

Figure S1. Distribution of RINT-transformed serum protein levels ....3

Figure S2. Associations with GRS64 ....5

Figure S3. Associations with MHC-only GRS ....7

Figure S4. Associations with non-MHC GRS ....9

Figure S5. RINT transformation distributions ....11

**Supplementary Tables**

Table S1. Clinical characteristics ....12

Table S2. Composite score associations ....13

Table S3. GRS correlations ....14

Table S4. Assay performance ....15

Table S5. Genetic variants and weights ....16

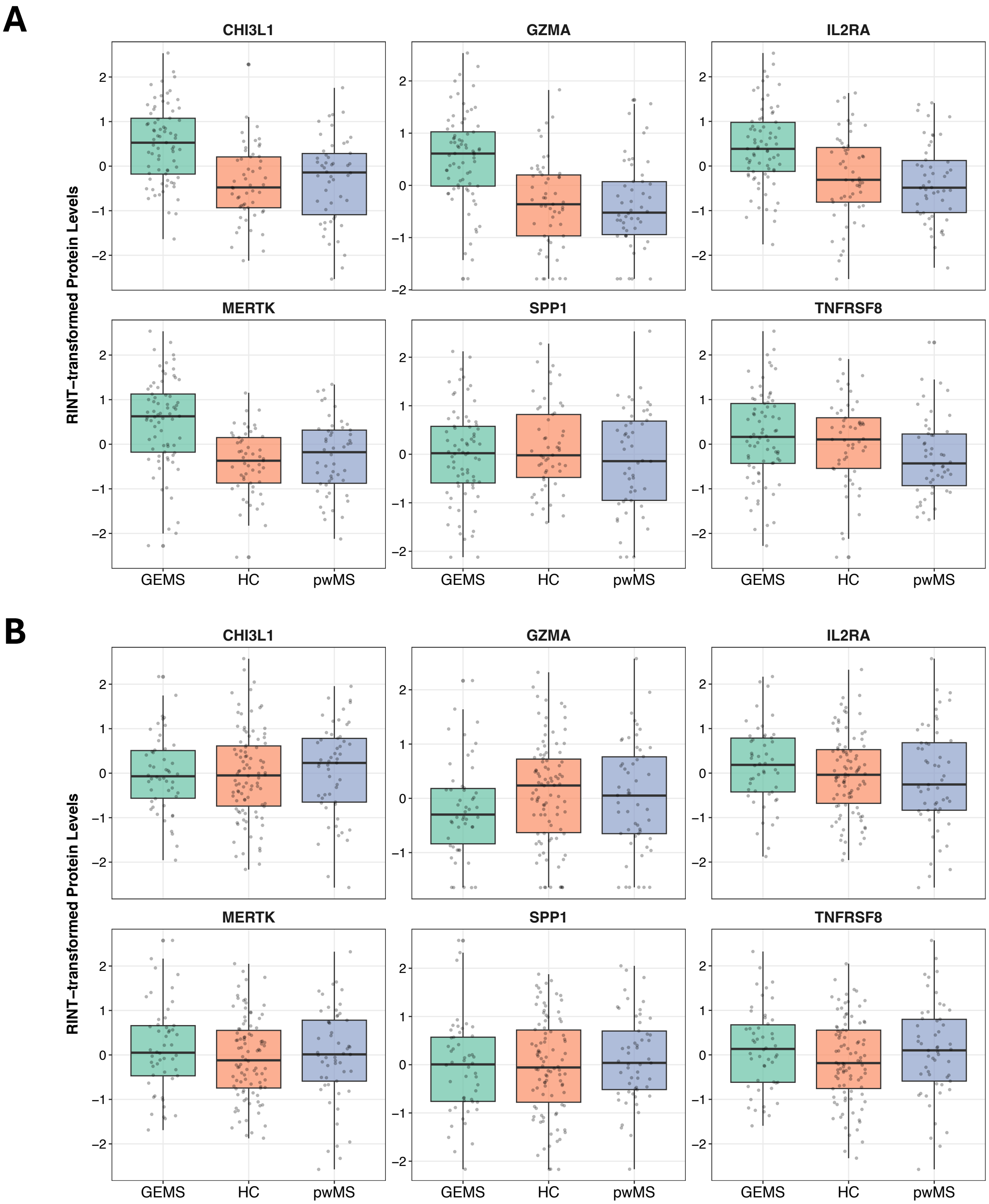


**Supplementary Figure 1. Distribution of RINT-transformed serum protein levels by study group.** Box plots show RINT-transformed protein levels in **(A)** CUIMC and **(B)** NINDS sample collections. Each box represents the interquartile range (IQR), with the horizontal line indicating the median; whiskers extend to 1.5× the IQR, and points represent individual participants. Group differences between pwMS (orange), healthy control participants (HC) (green), and GEMS participants (blue) were assessed using linear regression adjusted for age, sex, BMI, and batch, with Benjamini–Hochberg false discovery rate (FDR) correction for multiple testing.

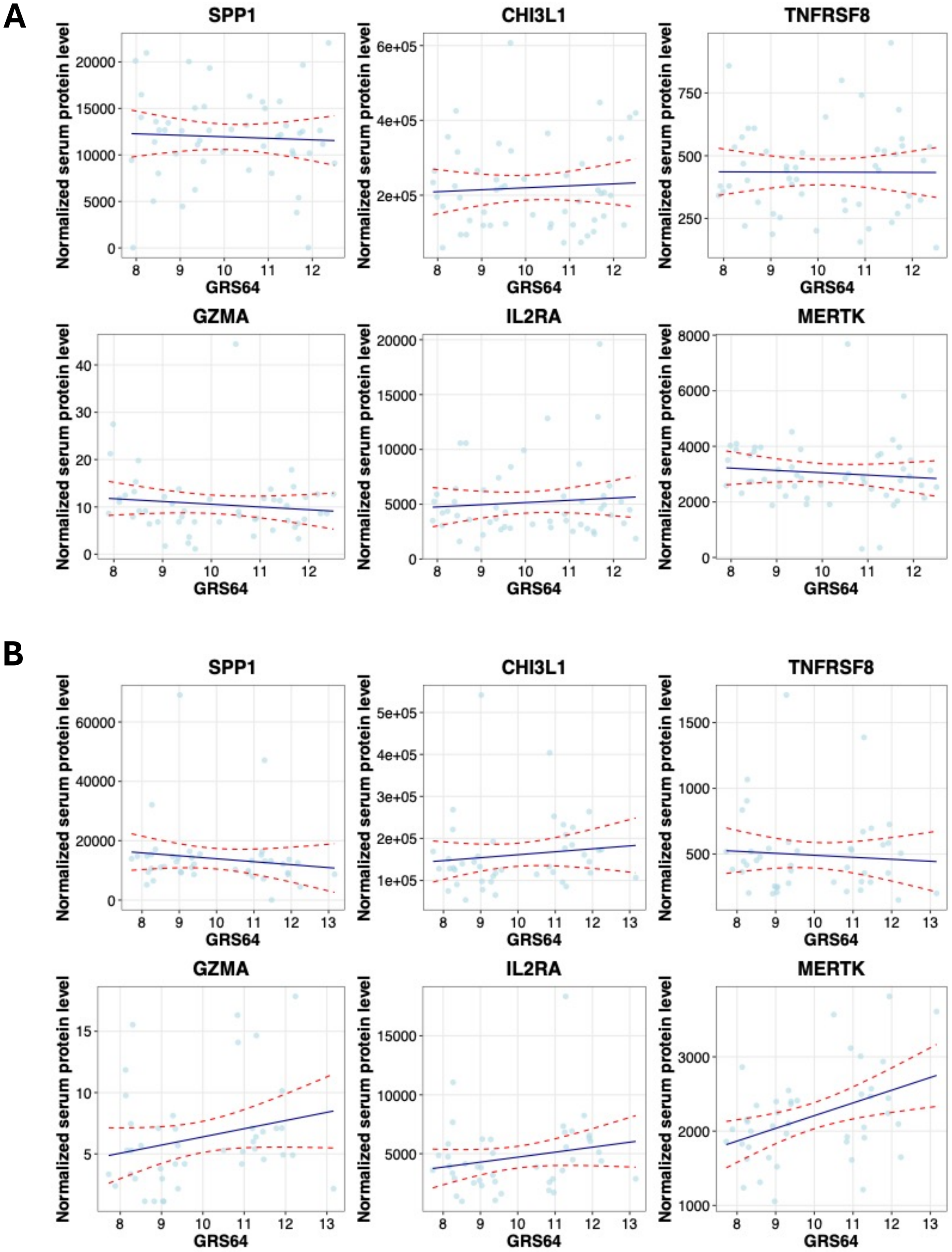


**Supplementary Figure 2. Associations between serum protein levels and MS GRS64.** Scatter plots show the relationship between the indicated genetic risk score and RINT-transformed serum protein levels in **(A)** CUIMC and **(B)** NINDS sample collections. Each point represents an individual participant. Lines indicate linear regression fits with 95% confidence intervals.

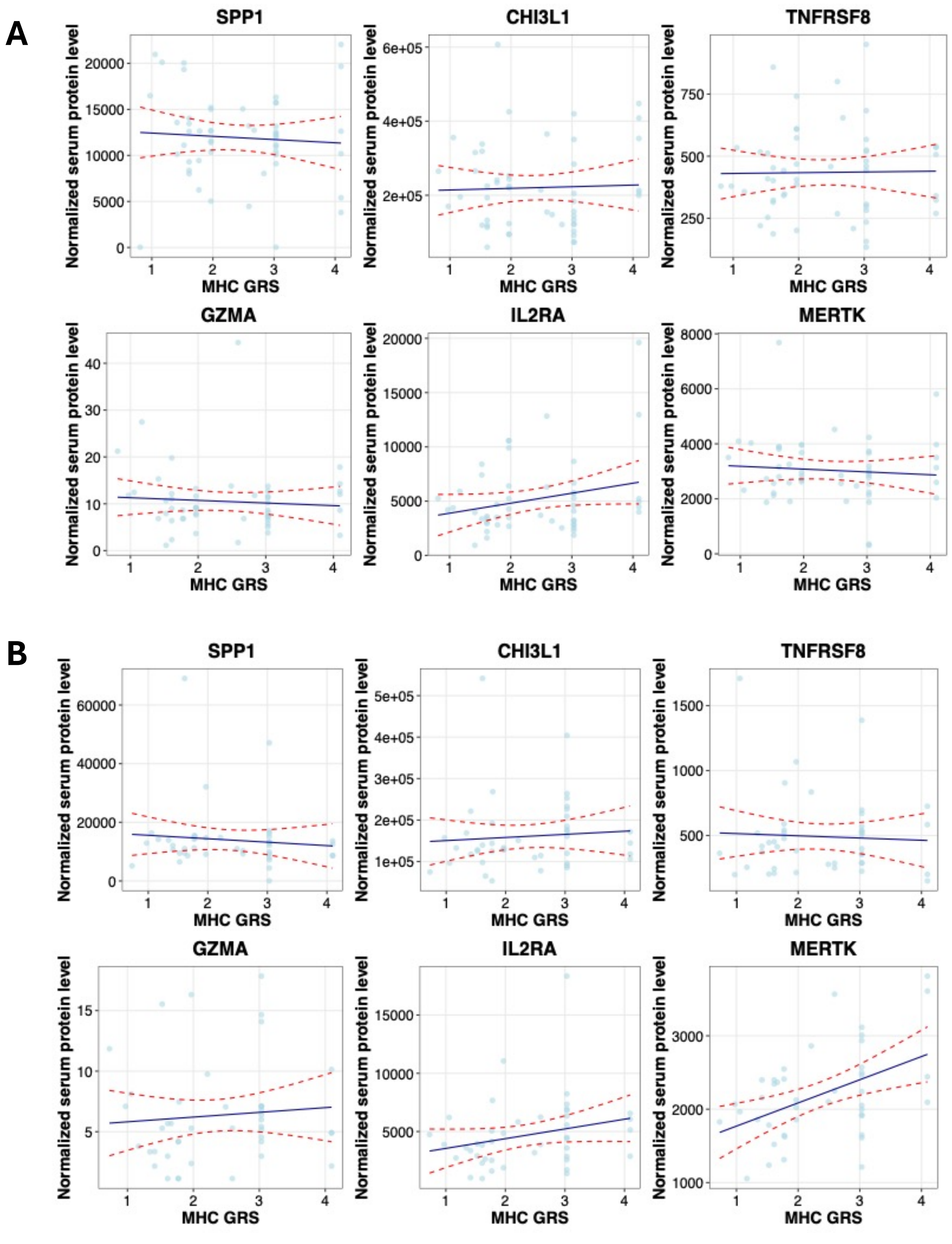


**Supplementary Figure 3. Associations between serum protein levels and MS MHC-only GRS.** Scatter plots show the relationship between the indicated genetic risk score and RINT-transformed serum protein levels in **(A)** CUIMC and **(B)** NINDS sample collections. Each point represents an individual participant. Lines indicate linear regression fits with 95% confidence intervals.

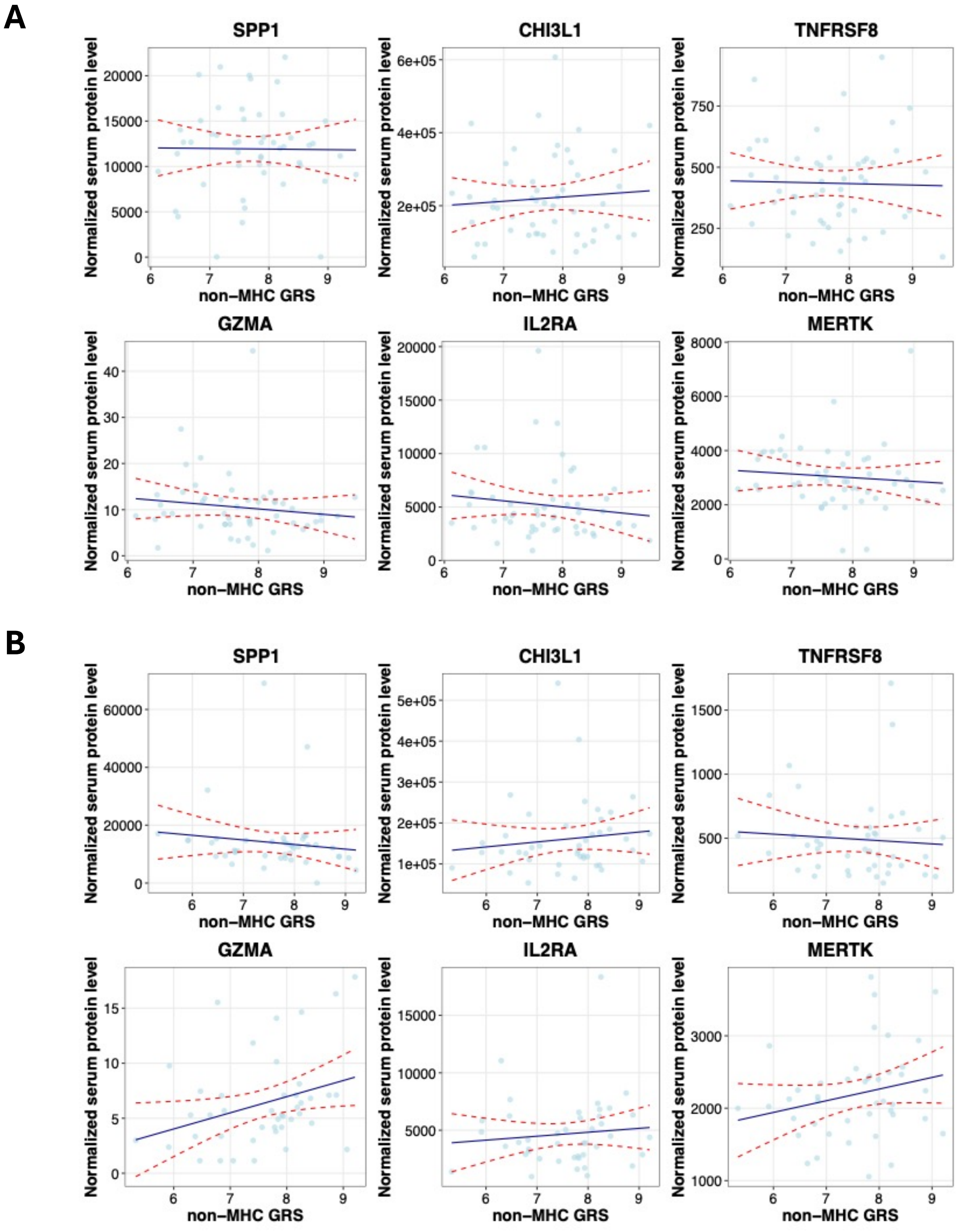


**Supplementary Figure 4. Associations between serum protein levels and non-MHC MS GRS.** Scatter plots show the relationship between the indicated genetic risk score and RINT-transformed serum protein levels in **(A)** CUIMC and **(B)** NINDS sample collections. Each point represents an individual participant. Lines indicate linear regression fits with 95% confidence intervals.

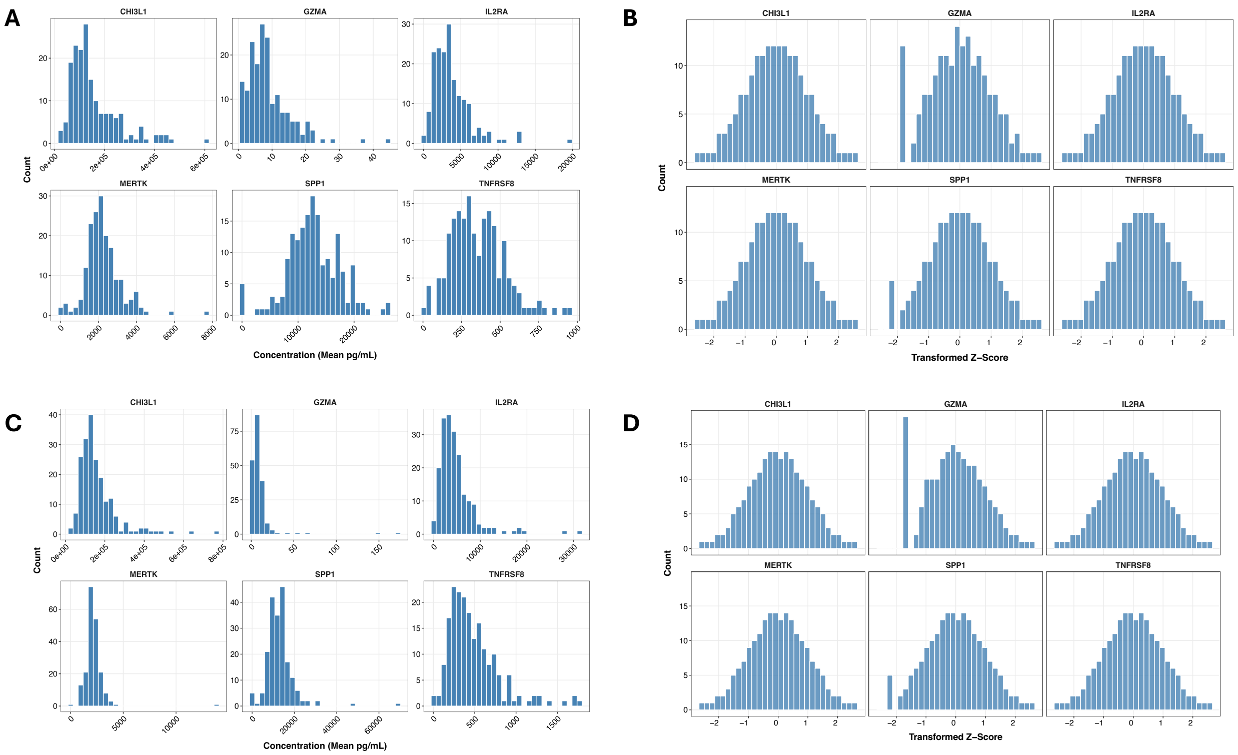


**Supplementary Figure 5. Rank-inverse normal transformation of serum protein measurements.** Histograms illustrating the distribution of protein measurements before **(A,** CUIMC**; C,** NINDS**)** and after **(B,** CUIMC**; D,** NINDS**)** rank-inverse normal transformation (RINT). Each panel displays the six analytes included in the multiplex panel (TNFRSF8, GZMA, IL2RA, MERTK, SPP1, and CHI3L1).

| Variable | CUIMC pwMS (n = 50) | NINDS pwMS (n = 53) |
|---|---|---|
| **Diagnosis (n)** | | |
| RRMS | 36 | 16 |
| CIS | 4 | 20 |
| RIS | 3 | 3 |
| SPMS | 4 | 2 |
| PPMS | 3 | 12 |
| **Therapy (none / steroids / DMT) (n)** | 32 / 5 / 13 | 50 / 0 / 3 |

**Supplementary Table 1. Clinical characteristics of MS participants in the CUIMC and NINDS sample collections.** Counts of MS diagnostic subtypes and treatment status at the time of serum collection. Diagnostic categories include relapsing–remitting MS (RRMS), clinically isolated syndrome (CIS), radiologically isolated syndrome (RIS), secondary progressive MS (SPMS), and primary progressive MS (PPMS). Treatment status reflects the number of participants receiving no therapy, corticosteroids, or disease-modifying therapies (DMTs) at the time of serum collection. Differences in subtype distribution between collections reflect differences in recruitment and clinical context; CUIMC participants were recruited during diagnostic evaluation, whereas NINDS samples were obtained cross-sectionally as part of a natural history protocol.

| | CUIMC | | | | NINDS | | | |
|---|---|---|---|---|---|---|---|---|
| | Estimate | SE | t value | p value | Estimate | SE | t value | p value |
| pwMS* | -74.102 | 100.583 | -0.737 | 0.463 | 127.046 | 157.816 | 0.805 | 0.422 |
| GEMS** | – | – | – | – | 277.724 | 155.591 | 1.785 | 0.076 |
| Age*** | -4.404 | 4.232 | -1.041 | 0.301 | -0.483 | 5.106 | -0.095 | 0.925 |
| Male Sex*** | 26.245 | 102.526 | 0.256 | 0.798 | -10.253 | 128.788 | -0.08 | 0.937 |

*Results compare pwMS and HC within each collection.
**Results for GEMS are calculated only for the NINDS collection, relative to HC; CUIMC GEMS were excluded due to sample collection differences.
***Covariate associations were estimated within each collection using the same multivariable models; across all included participants (CUIMC: HC and pwMS; NINDS: HC, pwMS, and GEMS), with adjustment for BMI, batch, and study group, such that estimates reflect associations independent of subgroup origin within collection.

**Supplementary Table 2. Association between the serum composite protein score and study groups.** The composite score was constructed as a weighted combination of six serum proteins (GZMA, MERTK, IL2RA, SPP1, TNFRSF8, and CHI3L1) using previously reported effect sizes[12]. Individual protein measurements were rank-inverse normal transformed (RINT) prior to calculation of the composite score. Linear regression models were fit separately for each collection (CUIMC and NINDS) using the composite score as the outcome. For group comparisons, models included study group (Subject_Type) and were adjusted for age, sex, BMI, and experimental date/batch. Values are reported as regression coefficients with standard errors (β [SE]) and corresponding p-values.

| | GRS metric | CUIMC r | CUIMC p | CUIMC FDR | NINDS r | NINDS p | NINDS FDR |
|---|---|---|---|---|---|---|---|
| GZMA | MHC | -0.075 | 0.603 | 0.916 | 0.084 | 0.590 | 0.749 |
| | non-MHC | -0.135 | 0.344 | 0.875 | 0.318 | 0.038 | 0.227 |
| | GRS64 | -0.121 | 0.396 | 0.883 | 0.241 | 0.120 | 0.346 |
| MERTK | MHC | -0.081 | 0.574 | 0.916 | 0.463 | 0.002 | 0.011 |
| | non-MHC | -0.093 | 0.516 | 0.875 | 0.236 | 0.128 | 0.383 |
| | GRS64 | -0.102 | 0.476 | 0.883 | 0.419 | 0.005 | 0.031 |
| IL2RA | MHC | 0.247 | 0.081 | 0.486 | 0.251 | 0.105 | 0.315 |
| | non-MHC | -0.131 | 0.358 | 0.875 | 0.103 | 0.511 | 0.614 |
| | GRS64 | 0.085 | 0.555 | 0.883 | 0.212 | 0.173 | 0.346 |
| SPP1 | MHC | -0.067 | 0.642 | 0.916 | -0.096 | 0.542 | 0.749 |
| | non-MHC | -0.011 | 0.939 | 0.939 | -0.130 | 0.406 | 0.614 |
| | GRS64 | -0.048 | 0.736 | 0.883 | -0.135 | 0.388 | 0.529 |
| TNFRSF8 | MHC | 0.015 | 0.916 | 0.916 | -0.050 | 0.751 | 0.751 |
| | non-MHC | -0.026 | 0.857 | 0.939 | -0.074 | 0.639 | 0.639 |
| | GRS64 | -0.005 | 0.974 | 0.974 | -0.074 | 0.637 | 0.637 |
| CHI3L1 | MHC | 0.035 | 0.810 | 0.916 | 0.077 | 0.624 | 0.749 |
| | non-MHC | 0.079 | 0.583 | 0.875 | 0.124 | 0.427 | 0.614 |
| | GRS64 | 0.065 | 0.650 | 0.883 | 0.121 | 0.441 | 0.529 |

**Supplementary Table 3. Correlation between serum protein levels and MS genetic risk scores in GEMS participants.** Within each collection, Pearson correlation coefficients were calculated between RINT-transformed protein levels and three MS genetic risk scores: (1) GRS64, aggregating the effects of 64 MS-associated variants; (2) an MHC-specific GRS, reflecting risk attributable to variants within the major histocompatibility complex (MHC); and (3) a non-MHC GRS, encompassing the cumulative contribution of non-MHC susceptibility loci. Correlation coefficients (r), nominal p values (p), and FDR values are reported separately for each collection. These analyses represent unadjusted associations.

| | Intra-Assay CV (%) | Inter-Assay CV (%) All Batches | Inter-Assay CV (%) Excluding Batch 1 |
|---|---|---|---|
| **TNFRSF8** | 7.86 | 7.1 | 6.57 |
| **GZMA** | 21.18 | 23.97 | 25.4 |
| **IL2RA** | 8.2 | 7.59 | 7.09 |
| **MERTK** | 7.09 | 8.88 | 9.69 |
| **SPP1** | 13.11 | 6.48 | 6.85 |
| **CHI3L1** | 9.02 | 11.79 | 12.17 |

**Supplementary Table 4. Assay Performance.** Intra-assay coefficients of variation (CV) were calculated from duplicate wells measured within the same assay plate. Inter-assay CVs were calculated using pooled serum control samples included across assay runs. In the first assay batch, the pooled control sample was prepared fresh, whereas frozen aliquots of the same pool were used in subsequent batches. Because this difference may influence inter-assay variability, CVs are reported both including all assay batches and after exclusion of the first batch. CV values are reported as percentages for each analyte.

| SNP (rsID) | Chromosome | Gene / Locus | Risk Allele | Alternative Allele | Risk Allele Frequency | Weight (log OR) | % Total GRS | MHC Region | Reference |
|---|---|---|---|---|---|---|---|---|---|
| rs3129889 | 6 | HLA-DRB1*15:01 | G | A | 0.224 | 1.0639 | 0.1150 | Yes | 50 |
| rs2395175 | 6 | HLA-DRB1*04:01 | G | A | 0.909 | 0.4415 | 0.0477 | Yes | PC |
| rs2844821 | 6 | HLA-A*02:01 | T | C | 0.882 | 0.3577 | 0.0387 | Yes | PC |
| rs3817964 | 6 | HLA-DRB1*04:04 | A | T | 0.051 | 0.2445 | 0.0264 | Yes | PC |
| rs12722489 | 10 | IL2RA | C | T | 0.878 | 0.2184 | 0.0236 | No | 50 |
| rs2300747 | 1 | CD58 | A | G | 0.903 | 0.2102 | 0.0227 | No | 50 |
| rs2119704 | 14 | GPR65 | C | A | 0.947 | 0.2095 | 0.0226 | No | 49 |
| rs4613763 | 5 | PTGER4 | C | T | 0.139 | 0.1964 | 0.0212 | No | 49 |
| rs498422 | 6 | HLA-DRB1*14:01 | T | G | 0.958 | 0.1847 | 0.0200 | Yes | PC |
| rs7089861 | 10 | IL2RA | C | G | 0.882 | 0.1806 | 0.0195 | No | 50 |
| rs2293152 | 17 | STAT3 | C | G | 0.652 | 0.1693 | 0.0183 | No | 50 |
| rs7200786 | 16 | CLEC16A | A | G | 0.487 | 0.1654 | 0.0179 | No | 49 |
| rs1077667 | 19 | TNFSF14 | C | T | 0.832 | 0.1519 | 0.0164 | No | 49 |
| rs4648356 | 1 | MMEL1 | C | A | 0.700 | 0.1475 | 0.0159 | No | 49 |
| rs10201872 | 2 | SP140 | T | C | 0.187 | 0.1391 | 0.0150 | No | 49 |
| rs170934 | 3 | EOMES | T | C | 0.501 | 0.1360 | 0.0147 | No | 50 |
| rs17066096 | 6 | IL22RA2 | G | A | 0.258 | 0.1355 | 0.0146 | No | 49 |
| rs874628 | 19 | MPV17L2 | A | G | 0.739 | 0.1343 | 0.0145 | No | 49 |
| rs1132200 | 3 | TMEM39A | C | T | 0.872 | 0.1330 | 0.0144 | No | 50 |
| rs6074022 | 20 | CD40 | C | T | 0.275 | 0.1329 | 0.0144 | No | 50 |
| rs17824933 | 11 | CD6 | G | C | 0.242 | 0.1314 | 0.0142 | No | 50 |
| rs1800693 | 12 | TNFRSF1A | C | T | 0.428 | 0.1269 | 0.0137 | No | 50 |
| rs2760524 | 1 | RGS1 | G | A | 0.856 | 0.1237 | 0.0134 | No | 50 |
| rs650258 | 11 | CD5 | C | T | 0.671 | 0.1228 | 0.0133 | No | 49 |
| rs13192841 | 6 | NA | A | G | 0.298 | 0.1217 | 0.0132 | No | 49 |
| rs12466022 | 2 | NA | C | A | 0.739 | 0.1210 | 0.0131 | No | 49 |
| rs11154801 | 6 | AHI1 | A | C | 0.368 | 0.1197 | 0.0129 | No | 49 |
| rs2303759 | 19 | DKKL1 | G | T | 0.279 | 0.1186 | 0.0128 | No | 49 |
| rs703842 | 12 | METTL1 | A | G | 0.714 | 0.1154 | 0.0125 | No | 50 |
| rs1250550 | 10 | ZMIZ1 | A | C | 0.329 | 0.1143 | 0.0124 | No | 49 |
| rs744166 | 17 | STAT3 | G | A | 0.441 | 0.1147 | 0.0124 | No | 50 |
| rs1738074 | 6 | TAGAP | C | T | 0.705 | 0.1137 | 0.0123 | No | 49 |
| rs802734 | 6 | PTPRK | A | G | 0.734 | 0.1126 | 0.0122 | No | 49 |
| rs6897932 | 5 | IL7R | C | T | 0.770 | 0.1119 | 0.0121 | No | 50 |
| rs17445836 | 16 | IRF8 | G | A | 0.814 | 0.1120 | 0.0121 | No | 50 |
| rs2744148 | 16 | SOX8 | G | A | 0.203 | 0.1107 | 0.0120 | No | 49 |
| rs12122721 | 1 | KIF21B | G | A | 0.737 | 0.1081 | 0.0117 | No | 50 |
| rs354033 | 7 | ZNF767 | G | A | 0.752 | 0.1080 | 0.0117 | No | 49 |
| rs11129295 | 3 | EOMES | T | C | 0.392 | 0.1078 | 0.0117 | No | 49 |
| rs11581062 | 1 | SLC30A7 | G | A | 0.317 | 0.1075 | 0.0116 | No | 49 |
| rs12048904 | 1 | EXTL2 | T | C | 0.354 | 0.1056 | 0.0114 | No | 49 |
| rs17174870 | 2 | MERTK | C | T | 0.777 | 0.1037 | 0.0112 | No | 49 |
| rs4680534 | 3 | IL12A | C | T | 0.362 | 0.1037 | 0.0112 | No | 50 |
| rs1790100 | 12 | MPHOSPH9 | G | T | 0.251 | 0.1034 | 0.0112 | No | 50 |
| rs4902647 | 14 | ZFP36L1 | C | T | 0.540 | 0.1031 | 0.0111 | No | 49 |
| rs2019960 | 8 | PVT1 | C | T | 0.217 | 0.1027 | 0.0111 | No | 49 |
| rs4410871 | 8 | MYC | C | T | 0.741 | 0.1018 | 0.0110 | No | 49 |
| rs13333054 | 16 | IRF8 | T | C | 0.233 | 0.1019 | 0.0110 | No | 49 |
| rs180515 | 17 | RPS6KB1 | G | A | 0.376 | 0.1005 | 0.0109 | No | 49 |
| rs12368653 | 12 | AGAP2 | A | G | 0.526 | 0.0970 | 0.0105 | No | 49 |
| rs771767 | 3 | NFKBIZ | A | G | 0.255 | 0.0963 | 0.0104 | No | 49 |
| rs4285028 | 3 | SLC15A2 | A | C | 0.740 | 0.0952 | 0.0103 | No | 49 |
| rs2300603 | 14 | BATF | T | C | 0.776 | 0.0933 | 0.0101 | No | 49 |
| rs12212193 | 6 | BACH2 | G | A | 0.461 | 0.0889 | 0.0096 | No | 49 |
| rs8112449 | 19 | TYK2, CDC37 | G | A | 0.687 | 0.0894 | 0.0097 | No | 49, PC |
| rs6718520 | 2 | Intergenic | A | G | 0.459 | 0.0852 | 0.0092 | No | 50 |
| rs7090512 | 10 | IL2RA | C | T | 0.360 | 0.0843 | 0.0091 | No | 49 |
| rs9846534 | 3 | CBLB | T | C | 0.825 | 0.0765 | 0.0083 | No | 50 |
| rs4308217 | 3 | CD86 | C | A | 0.687 | 0.0745 | 0.0080 | No | 49 |
| rs1393122 | 5 | Intergenic | A | G | 0.857 | 0.0619 | 0.0067 | No | 50 |
| rs2762932 | 20 | CYP24A1 | C | T | 0.153 | 0.0589 | 0.0064 | No | 49 |
| rs763361 | 18 | CD226 | T | C | 0.494 | 0.0439 | 0.0047 | No | 50 |

**Abbreviations:** SNP, single nucleotide polymorphism; MHC, major histocompatibility complex; PC, personal communication from International Multiple Sclerosis Genetics Consortium

**a** Risk allele frequency reflects the observed frequency among genotyped participants in the GEMS cohort.

**b** Effect sizes represent the natural logarithm of published odds ratios for multiple sclerosis susceptibility and were used as weights in construction of the genetic risk score.

**c** % Total GRS indicates the proportional contribution of each variant to the total genetic risk score based on its relative weight.

**d** Variants located within the MHC region include established HLA alleles (e.g., *HLA-DRB1**15:01, *HLA-A**02:01, *HLA-DRB1**04:01, *HLA-DRB1**04:04, and *HLA-DRB1**14:01).

**Supplementary Table 5. Genetic variants and weights used to construct the multiple sclerosis genetic risk score (GRS64).** The GRS64 was constructed using 64 previously identified MS susceptibility variants derived from published genome-wide association studies[49,50]. Each variant was coded additively according to the number of risk alleles and weighted by the natural logarithm of the reported odds ratio for MS susceptibility. Variants are shown with their corresponding genomic location, risk allele, allele frequency in the GEMS cohort, and contribution to the overall genetic risk score. Variants located within the major histocompatibility complex (MHC) region are indicated. Rows are ordered by descending effect size (weight).
