## Supplementary Methods for "Evaluation of potential serum biomarkers for individuals at risk of multiple sclerosis"

Additional methodological details supporting the primary analyses are provided below.

### Multiplex bead-based immunoassays

Serum protein concentrations were measured using custom multiplex bead-based immunoassays (ProcartaPlex, Thermo Fisher Scientific): a 4-plex assay (catalog number PPX-04-MXH6DA4) measuring TNFRSF8, MERTK, GZMA, and IL2RA, and a 2-plex assay (catalog number PPX-02-MXGZHP7) measuring SPP1 and CHI3L1 (assayed separately as they required a 1:100 dilution).

Immunoassays were performed according to the manufacturer's instructions and using the ProcartaPlex Platinum Assay Buffer (catalog number EPXP-11113-000) as per manufacturer's recommendations. Fluorescence intensity was measured using a Luminex 200 instrument (software xPONENT 4.2). Background signal was assessed using designated wells on each plate.

### Assay Performance Evaluation

A pooled serum bridging sample was included on each plate to monitor assay performance. Inter-assay variability was assessed using the bridging sample by calculating the mean concentration, standard deviation (SD), and coefficient of variation (%CV) across assay batches for each biomarker. Intra-assay variability was evaluated using duplicate wells for each sample, with mean concentration, SD, and %CV calculated from duplicate measurements. Because the bridging pool was freshly prepared for the first batch and frozen for subsequent batches, inter-assay variability was evaluated both including and excluding the first experimental run. Excluding the first experimental run, inter-assay CVs were generally low, ranging from 6.57% to 12.17% for five of the six analytes (**Supplementary Table 4**), while GZMA showed higher inter-assay variability (25.40%). Results were similar when the first run was included in the analysis. Intra-assay precision was similar, with the mean intra-assay CVs ranging from 7.09% to 13.11% (**Supplementary Table 4**), and GZMA demonstrated higher variability (21.18%). Overall, assay performance fell within the range of commonly accepted performance thresholds for multiplex bead-based immunoassays (intra-assay CV < 15% and inter-assay CV < 20%). Five of the six analytes met these criteria, while GZMA showed higher variability consistent with its relatively low circulating abundance[49].

**Software**

Data processing and visualization were conducted using packages from the tidyverse ecosystem (including dplyr, tidyr, and stringr), while statistical modeling and meta-analysis were performed using broom, car, caret, and metafor. Figures were generated using ggplot2 and related visualization packages (ggrepel, ggfortify, ggpubr), and data tables were exported using openxlsx.
